# Potassium-Selective Nanoelectrode Arrays for Single-Cell Profiling of human iPSC-Derived Cardiomyocytes

**DOI:** 10.64898/2026.02.13.705576

**Authors:** Dhivya Pushpa Meganathan, Romeo Banzon, Ana Casanova, Einollah Sarikhani, Kuldeep Mahato, Hillary Vu, Sarah Reade, Iswerya Ambika Devarajan, Anum Tahir, Lekshmi Sasi, Leah Sadr, Joseph Wang, Zeinab Jahed

## Abstract

Potassium ion (K⁺) dynamics are central to cardiac electrophysiology, with early disruptions in K⁺ flux often preceding arrhythmia and contractile dysfunction. However, current sensing technologies, such as patch-clamp, Microelectrode arrays (MEAs), and fluorescent indicators, either lack chemical specificity for K⁺ or are unsuitable for long-term, single-cell analysis. Conventional ion-selective electrodes (ISEs), while more selective, are limited by bulk-phase design and poor spatial resolution. To address these limitations, we present KINESIS (K⁺-Ion Nano-Electrode Selective Interface System), a nanofabricated, cell-compliant platform that enables direct, label-free potentiometric measurement of K⁺ gradients with subcellular precision. KINESIS features high-aspect-ratio nanopillars coated with a valinomycin-based K⁺ recognition membrane, forming a stable, non-invasive interface with human induced pluripotent stem cell-derived cardiomyocytes (hiPSC-CMs). This architecture allows localized, Nernstian sensing of K⁺ efflux or depletion without disrupting cell membranes. Pharmacological validation shows distinct potential shifts in response to caffeine and ouabain. KINESIS thus offers a highly selective, spatially resolved approach for studying K⁺ handling in cardiotoxicity screening and patient-specific disease modeling.

---

Nanopillar electrode arrays (NEAs) offer a powerful platform for high-throughput, parallel, single-cell recording of intracellular activity. Their vertical geometry enables minimally invasive access to subcellular space, forming intimate interfaces with the plasma membrane and allowing spatially resolved electrical recordings^1–3^. These features position NEAs as a compelling tool for large-scale electrophysiological measurements. However, existing NEA platforms lack ion selectivity and detect only aggregate voltage changes arising from the combined flux of multiple ions, including K⁺, Na⁺, Ca²⁺, and Cl⁻^1–6^. This lack of chemical resolution restricts their ability to isolate specific ionic events, limiting mechanistic interpretation of recorded signals. Targeting potassium is especially critical, as K⁺ dynamics governs membrane repolarization and directly influences excitability and rhythm^7^. Subtle shifts in K⁺ levels often precede arrhythmogenic triggers and serve as early markers of cardiac dysfunction^8–10^.

This limitation becomes especially problematic in contexts where distinguishing the behavior of individual ions is critical, such as drug screening, cardiotoxicity profiling, and disease modeling. Many pharmacological agents, such as cardiac glycosides like ouabain and digoxin, and methylxanthines like caffeine and theophylline, selectively target specific ion transporters or channels, and their effects cannot be resolved when measurements reflect mixed-ion signals^11–13^. There is a pressing need for bioelectronic platforms that combine the spatial resolution of NEAs with selective detection of individual ions like potassium. NEAs provide a scalable and minimally invasive scaffold ideally suited for hosting ion-selective coatings while preserving high-resolution, single-cell access.

Several classes of sensing technologies have attempted to address this need. These include (1) ion-selective electrodes (ISEs), (2) solid-state sensing platforms such as ion-sensitive field-effect transistors (ISFETs) ^14^, (3) optical imaging techniques using either dyes or genetically encoded sensors, and (4) vibrational spectroscopic approaches such as Raman-based methods. For enhancing chemical selectivity, ISEs have been developed to target specific ionic species^15^. ISEs are capable of highly selective detection through the use of ionophore receptors such as valinomycin, which provide strong affinity for K^+^. Despite their selectivity, traditional ISEs are generally designed for bulk-phase measurements and are ill-suited for integration with single-cell systems^16,17^. Their large size and rigid configuration hinder conformal contact with soft, dynamic cell membranes, and their operation under diffusion-limited, zero-current steady-state conditions constrains their ability to capture fast, localized ionic fluctuations. Solid-state sensing platforms, particularly as *ion-sensitive* field-effect transistor (ISFET)-based sensors, have emerged as promising tools for label-free, real-time detection of ionic activity. While ISFETs demonstrate chemical specificity when functionalized with ion-selective membranes and have shown success in extracellular ion monitoring, their adaptation to dynamic, single-cell electrophysiology remains constrained by limited surface proximity, long-term stability, and challenges in scaling for subcellular spatial resolution.^16,18^. Optical approaches, including small-molecule fluorescent probes (e.g., PBFI, APG-1) and genetically encoded potassium indicators (e.g., GEPIIs, GINKO)—provide spatial mapping of K^+^ flux but suffer from key limitations^19,20^. Fluorescent dyes often display limited selectivity over competing ions, low dynamic range, and poor photostability^21^. Genetically encoded sensors, although more selective, require stable gene delivery and expression, which poses significant challenges in primary human cells and iPSC-derived cardiomyocytes, particularly in clinical or high-throughput settings^19^. Furthermore, these sensors often perturb native cell function or require extensive optical instrumentation. Raman-based methods offer a label-free alternative for ionic detection but suffer from weak signal intensities, limited temporal resolution, and complex instrumentation, making them impractical for real-time, live-cell applications^22^. Collectively, these limitations underscore the critical need for a sensing platform capable of direct, ion-selective, non-invasive, and spatially localized detection of K^+^ dynamics in live cells.

To bridge nanoscale electrophysiology with chemical specificity, we developed KINESIS (K⁺-Ion Nano-Electrode Selective Interface System), the first nanoelectrode array platform capable of selective, real-time detection of potassium dynamics at the single-cell level. KINESIS integrates high-aspect-ratio platinum nanopillars with a valinomycin-based potassium-selective membrane, forming a mechanically stable, non-disruptive interface with the plasma membrane. This configuration enables localized transduction of potassium activity via the Nernst equation, supporting continuous, label-free sensing with subcellular resolution.

We validated KINESIS in human iPSC-derived cardiomyocytes by applying pharmacological agents with known effects on K^+^ handling. Caffeine, which enhances K^+^ efflux through calcium-activated SK channels, induced a characteristic positive potential shift. In contrast, ouabain, a Na^+^/K^+^-ATPase inhibitor, triggered a negative shift consistent with suppressed K^+^ reuptake.

These distinct signatures confirm the platform’s ability to resolve pharmacologically induced potassium dynamics in real time, including direction-specific changes in membrane-proximal K⁺ concentration consistent with known channel activity as shown in **Figure 1**.

**Figure 1.**
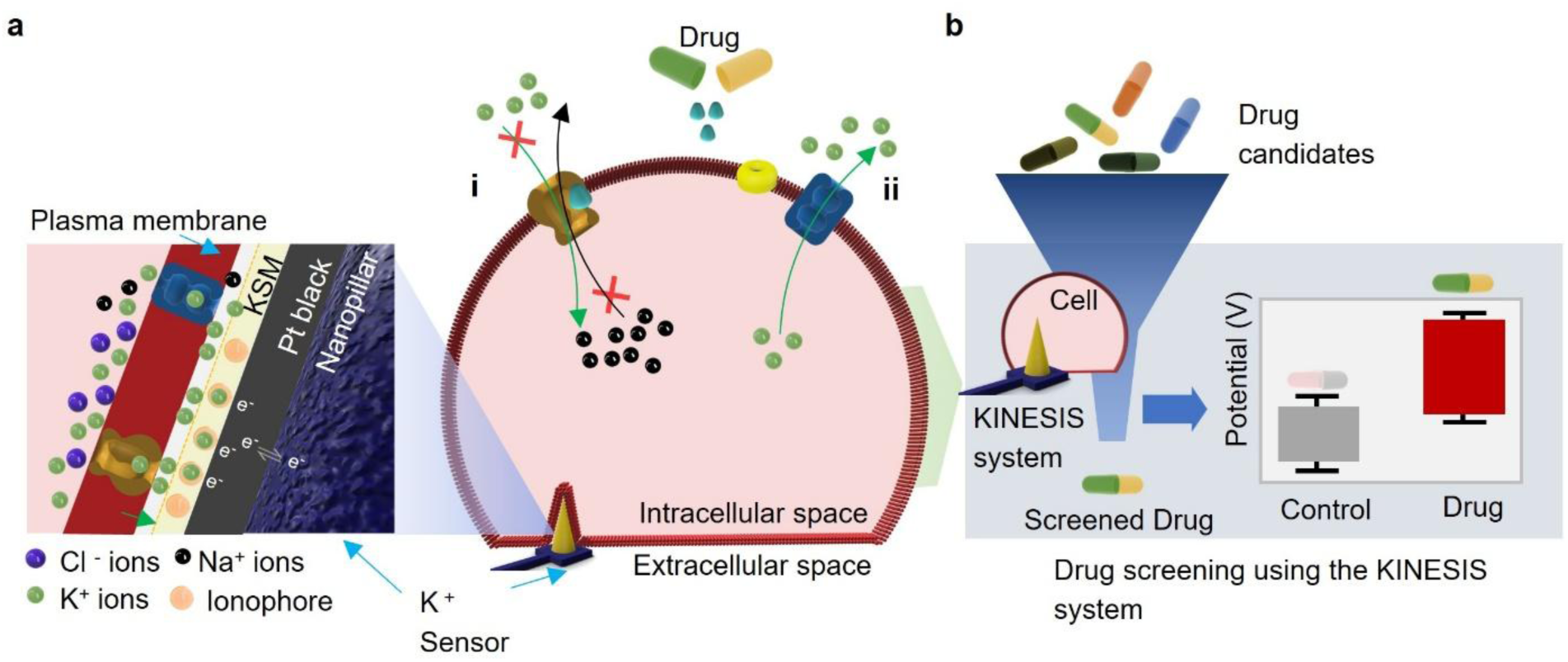
**Schematic representation of the development of the KINESIS system**: a. scheme illustrating the impact of the drug on the cardiomyocytes and the release of K^+^ ion to the extracellular space, thereby interacting to the nanopillar based solid-state K^+^ sensor. b. Strategy showing the novel drug screening system using the KINESIS platform.

By combining nanoscale access with chemical specificity, KINESIS overcomes critical limitations of current NEAs, ISEs, and optical sensors. Its modular, scalable design makes it a promising tool for integration into high-throughput drug screening pipelines, fundamental ion channel research, and personalized disease modeling in cardiovascular systems.

## RESULTS AND DISCUSSION

### Fabrication and characterization of the nanoelectrode array platform for parallel single-cell recording

Our sensing platform is built around a high-density array of 60 individually addressable platinum nanopillar electrodes. Scanning electron microscopy (SEM) images at increasing magnification **(Figure 2b-i to 2b-iii)** illustrates the array layout and electrode morphology. Each electrode consists of one to four free-standing tapered nanopillars (tip radius ∼200 nm, height: ∼7.2 µm, pitch: ∼12.4 *µ*m), fabricated by top-down etching into a fused silica substrate, following methods described previously. The total footprint of each nanoelectrode is 16 µm x 16 µm, which is well-matched to the spread area of a single hiPSC-CM **(Figure 2c)**, making the platform suitable for single-cell measurements^23^.

**Figure 2.**
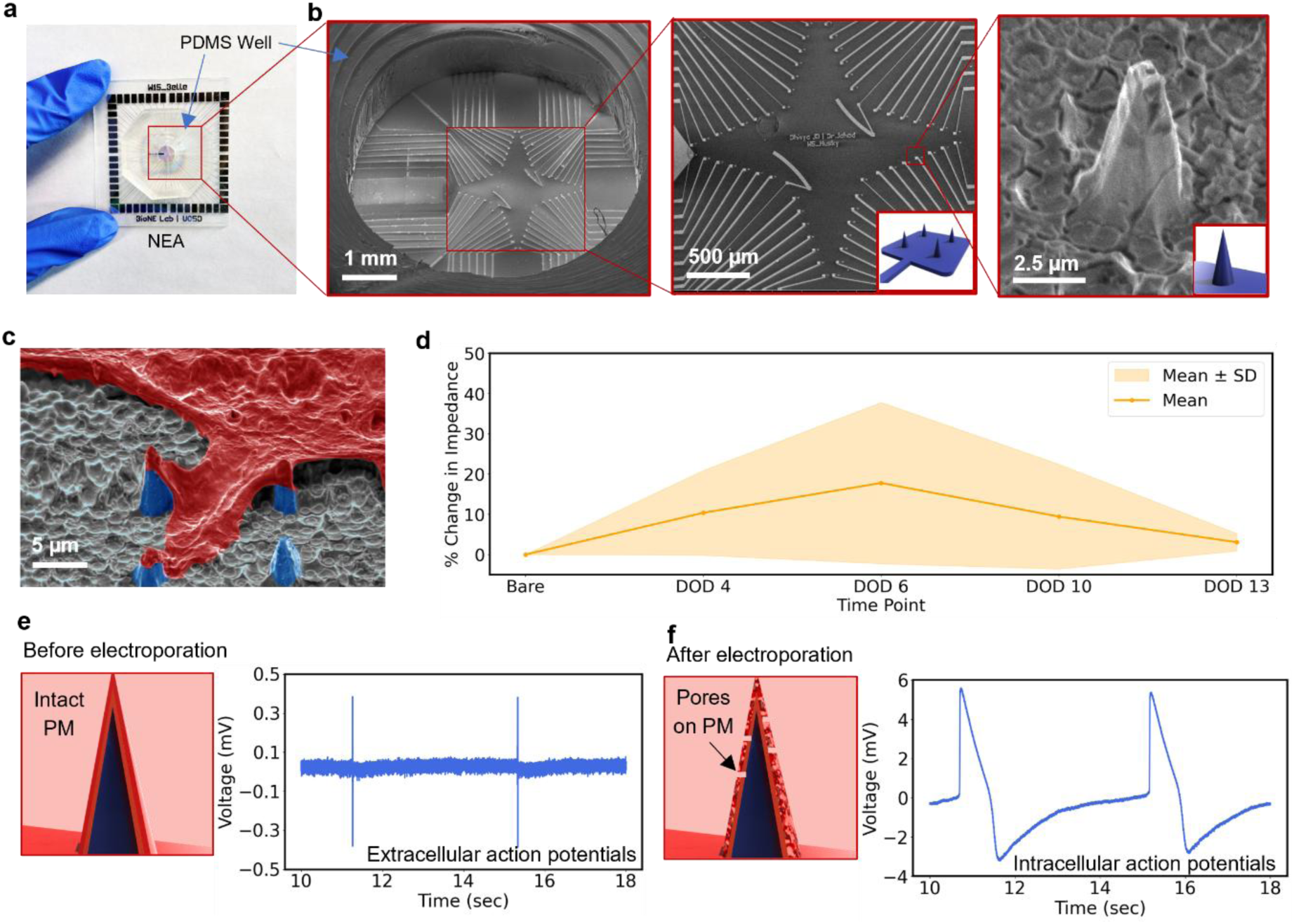
Characterization of Nanopillar Electrode Array: (a) Photographic image of the fabricated nanopillar electrode device. (b) SEM images of the nanopillar array: (i) Overview of the well (scale bar: 1mm); (ii) Zoomed-in view of the array (scale bar: 500µm); (iii) Single nanopillar (scale bar: 2.5µm). Insert showing schematic of the cell-nanoelectrode interface showing: (c) SEM image of the cell-nanoelectrode interface (scale bar: 5 µm) (d) Percent change in impedance at 1 kHz across Days on Devices (DOD) 4, 6, 10 and 13 relative to bare electrodes (e) Extracellular signal acquisition before electroporation; (f) Intracellular signal acquisition after electroporation showing pores created in Plasma membrane (PM).

To monitor the evolution of cell attachment and coupling, impedance at 1 kHz was recorded using the MEA-IT System (Multi Channel Systems). The instrument applies a 100 mV sinusoidal test signal and reports impedance magnitude at this frequency as a standard diagnostic measure of electrode contact quality^24^. The 1 kHz impedance increased by 10.37 ± 10.53 % on Day 4 and 17.79 ± 20.04 % on Day 6, indicating progressive cell–electrode sealing, and then decreased to 9.43 ± 13.02 % on Day 10 and 3.07 ± 2.15 % on Day 13, consistent with partial detachment or morphological changes (Figure 2d). These impedance trends, averaged across 60 electrodes under consistent conditions, were used qualitatively to identify Day 4 as the period of strongest and most stable cell–electrode coupling for subsequent recordings..

To further confirm and test electrode functionality, we performed electrical recordings of hiPSC-CMs. As shown in **Figure 2e and 2f**, the nanoelectrodes captured both extracellular and intracellular activity from spontaneously beating hiPSC-CMs, demonstrating reliable and high-fidelity signal acquisition at the single-cell level. Together, these findings confirm that our NEA platform enables robust, reproducible, and high-resolution interfacing with single hiPSC-CMs cells, an essential capability for intracellular electrophysiological measurements. Having established this functionality, we next focused on engineering chemically selective nanoelectrodes for targeted potassium ion sensing.

### Analytical performance assessment of KINESIS platform

To enable selective potassium detection on NEAs, we first screened Potassium ion-selective membrane (KSM) formulations using bulk Pt wire electrodes **(Figure S3a)**. This preliminary step allowed rapid benchmarking of membrane selectivity prior to NEA integration. Two formulations, KSM1 and KSM2, were tested. Both contained valinomycin, a K⁺-selective ionophore, but differed in ionophore-to-exchanger ratios: KSM1 used an excess of valinomycin (3.87:1), while KSM2 used equimolar amounts. To establish a baseline, we tested each membrane without ionophore. Both showed overlapping responses to KCl and NaCl, confirming minimal selectivity. Upon valinomycin incorporation, KSM1 produced a near-Nernstian slope (∼59 mV/decade) and effectively suppressed Na⁺ interference, whereas KSM2 remained less selective **(Figure S3c)**. This response is consistent with the established principle that maintaining an appropriate ionophore-to-ionic-site ratio ensures the presence of free (uncomplexed) valinomycin molecules for K⁺ exchange, preventing anionic interference and yielding a stable Nernstian behavior^25,26^. Based on its near-Nernstian slope and effective Na⁺ suppression, KSM1 was selected for subsequent testing on NEAs.t

Optimizing the transducer layer is critical for selective sensing because efficient ion-to-electron transduction improves potential stability, which reduces signal drift and ensures consistent selectivity under physiological conditions.. We next evaluated candidate transducer materials for their ability to support stable membrane adhesion, uniform coating, and functional electrical interfacing, using bulk Pt wire electrodes as a testing platform **(Figure S5a)**. Multiple conductive polymers and composites were first tested on Pt wires **(Figure S5a)**. Platinum Black (PtB) showed the strongest Nernstian response, consistent with its high surface area and enhanced charge transfer capacity^27,28^. Polyaniline doped with 1.5 equivalents of dodecylbenzenesulfonic acid (PANI:1.5DBSA) and Poly(3,4-ethylenedioxythiophene):poly(styrenesulfonate) ( PEDOT:PSS) also performed well (∼51 mV/decade), outperforming pristine PANI and graphene oxide (GO) composites^29^. Selectivity testing against Na^+^ **(Figure S5b)** revealed that PtB and PANI:1.5DBSA maintained K⁺ selectivity up to physiological sodium levels, while Polypyrrole (PPy) and GO-modified electrodes showed greater cross-reactivity. These results established PtB and PANI:1.5DBSA as viable transducer layers for NEA functionalization^30^.

Having identified optimal components in bulk tests, we next translated PtB and PANI:1.5DBSA onto NEAs **(Figure 3a)** for functional validation. Each material was electrodeposited onto separate nanopillar devices. To evaluate and compare their baseline electrical properties, we performed impedance spectroscopy prior to potassium-selective membrane (KSM1) application. PtB-modified NEAs exhibited significantly lower impedance at 1 kHz compared to PANI:1.5DBSA **(Figure 3c)**, consistent with efficient ion-to-electron transduction. SEM confirmed uniform membrane coverage **(Figure 3c)**, and the Schematic illustration of fully functionalized Nanopillar for K^+^ sensing and cross-sectional view of layers is shown in **Figure 3b**. Each device was coated with 2 µL of KSM1 and cured under ambient conditions for 24 hours to ensure membrane uniformity and adhesion.

**Figure 3.**
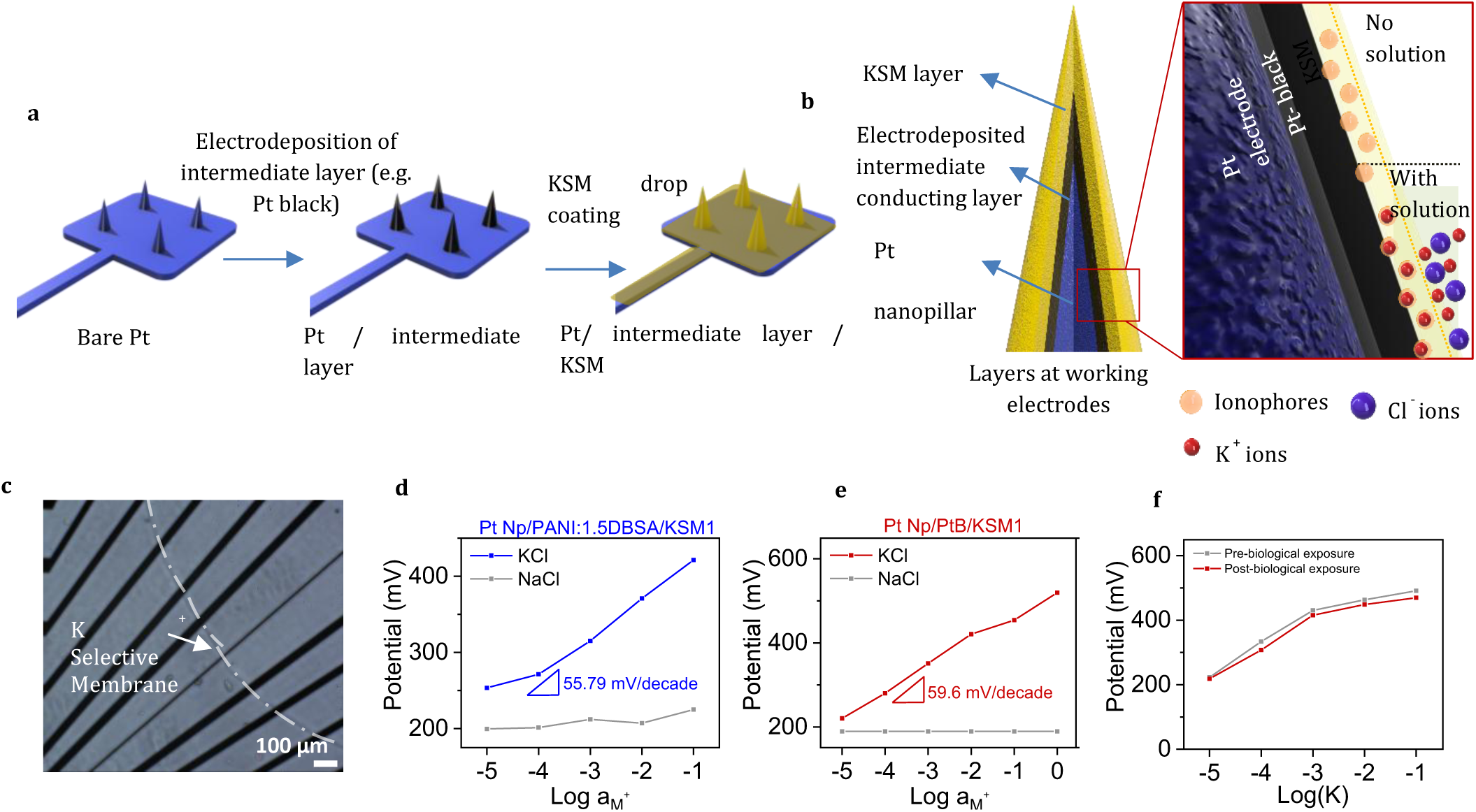
Functionalization of the Nanopillar Electrode Array (NEA) Platform and Membrane Characterization. (a) Schematic illustration of the KSM (potassium ion-selective membrane) coating process for functionalization of the NEA platform layers (b) Schematic illustration of fully functionalized Nanopillar for K+ sensing and cross-sectional view of layers (c) Microscopic image of the Potassium selective membrane trace on the device (scale bar 100 μm) Potassium calibration curves (potential vs. log[aM⁺] concentration): (d) PANI:1.5DBSA; (e) Pt Black, showing K+ sensitivity and Selectivity against NaCl interference, where aM⁺is the activity of the monovalent cation (f) Effect of biological exposure on K⁺ calibration. Potentiometric calibration curves (potential vs log[K⁺]) for a representative electrode measured before biological exposure (gray) and after two 5-day cardiomyocyte culture cycles on fibronectin-coated devices (red). The same electrode was recalibrated after each exposure and cleaning, with a total interval of approximately three months between the first and final calibration. Data for two additional electrodes showing similar trends are provided in Supplementary Figure S9.

We next evaluated sensor performance in the absence of cells via potentiometric calibration. Both configurations showed near-Nernstian slopes, 59.6 mV/decade for PtB and 55.8 mV/decade for PANI:1.5DBSA **(Figure 3d-e)**. At the transducer–membrane interface, PtB provided consistent ion-to-electron transduction with minimal potential variation, whereas PANI:1.5DBSA showed small baseline fluctuations at low K⁺ under the same conditions. These differences are consistent with established observations that metallic contacts offer chemically inert and stable charge-transfer behavior, while conducting polymer contacts, though effective, can exhibit slight potential shifts due to their redox-active nature and environmental sensitivity reported in prior SC-ISE studies^31^. Based on superior electrochemical performance, including lower impedance and a near-ideal Nernstian slope, PtB was selected for subsequent biological experiments. While PANI:1.5DBSA exhibited acceptable selectivity, small baseline variations at low K⁺ and a higher device impedance at 1 kHz prior to membrane application made it less suited for live-cell measurements. To confirm chemical selectivity under physiologically relevant ionic conditions, we recorded time-domain responses to sequential additions of K⁺, Ca²⁺, Mg²⁺, and Na⁺ (Supplementary Fig. S7g). Distinct potential steps were observed only for KCl, whereas Ca²⁺, Mg²⁺, and Na⁺ caused negligible changes, verifying that the valinomycin-based membrane maintains strong K⁺ specificity in the presence of competing mono- and divalent cations. Having established the device performance without cells, we next evaluated KINESIS in live-cell conditions. To further verify robustness and long-term stability under biological exposure, the same electrodes were recalibrated before and after a three-month interval during which they underwent two full cardiomyocyte culture and cleaning cycles. The representative electrode shown in Figure 3f retained stable near-Nernstian behavior after this period, showing only modest sensitivity loss (average slope change ≈ –12%) consistent with minor fouling or membrane aging. These results indicate that the valinomycin membrane preserved its ion-selective response despite repeated exposure to fibronectin coating, cell attachment, and protein-containing media, with comparable trends observed for additional electrodes shown in Supplementary Figure S9.

### KINESIS Enables Pharmacological Profiling of K⁺ Dynamics

Following functional validation on the NEA platform, we next assessed the biological performance of KINESIS by testing pharmacological models of potassium flux in live hiPSC-CMs. hiPSC-CMs were cultured directly on KINESIS as described in Methods. Photograph of 8 channel functionalized NEA device is shown in **Figure 4a**. Live and dead fluorescence images of iPSC-derived cardiomyocytes on the modified device (KINESIS) (Figure. 4b) showed a healthy, viable monolayer. Quantification of viability across both unmodified NEAs and Modified NEAs KINESIS (Figure. 4c) confirmed that the modification did not introduce cytotoxic effects, with no significant differences between the two conditions. Supplementary Video 1 further illustrates this monolayer confluency and spatial distribution across the platform. The vertical nanopillar geometry supports spatially confined cell–electrode interactions and may promote membrane wrapping, consistent with prior observations of curvature-induced deformation at nano-bio interfaces^32^. This close contact is expected to reduce cleft resistance and improve signal fidelity. Unlike electrophysiological approaches that record voltage transients, KINESIS transduces membrane-proximal K⁺ concentration into potential via a valinomycin-based ion-selective membrane, enabling subcellular chemical flux detection.

**Figure 4:**
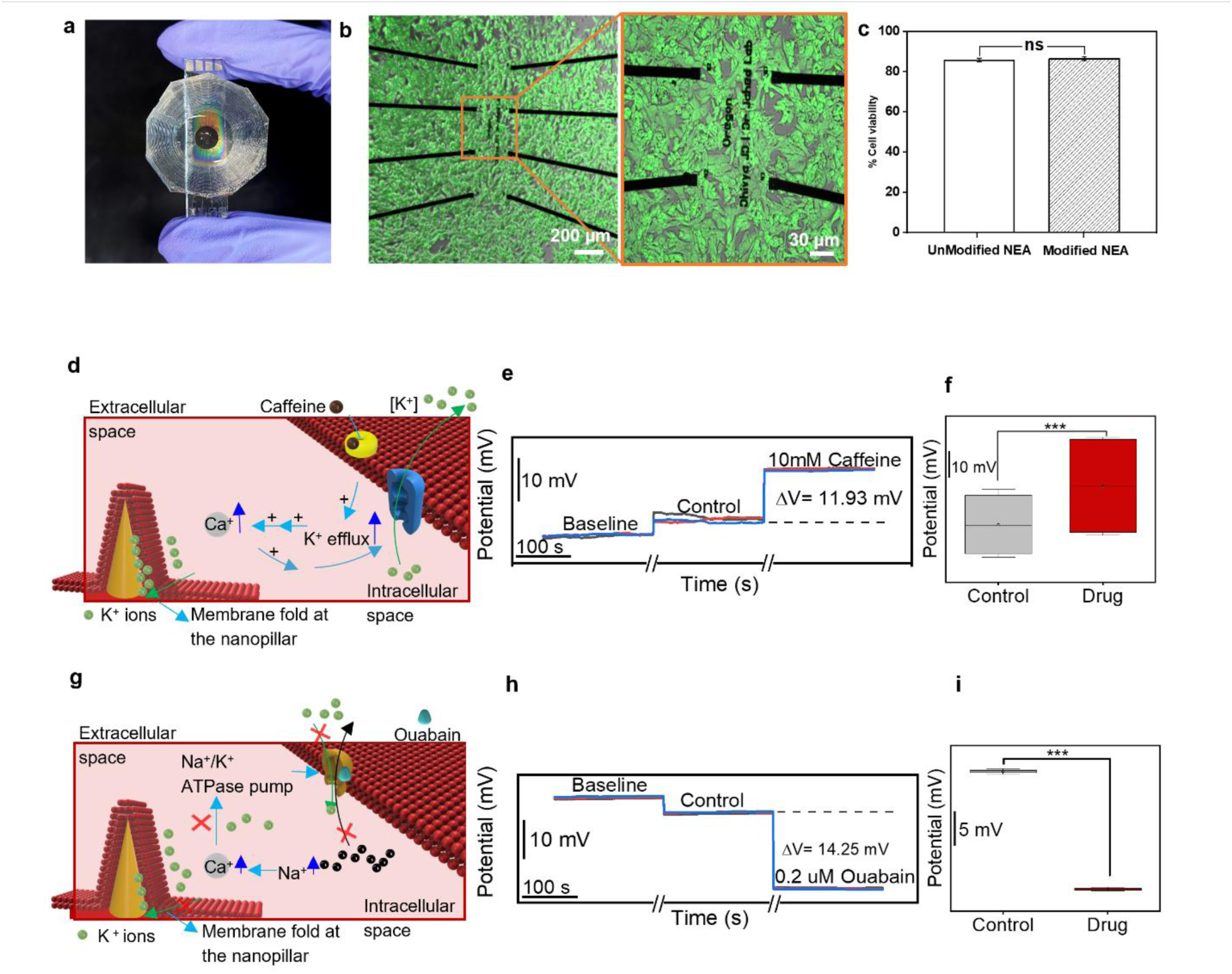
Validation of KINESIS platform for Measuring Drug-Induced Alterations in Cardiomyocyte Potassium Potentials. (a) Photograph of 8 channel functionalized NEA device (b) Live and dead assay fluorescence images of iPSC derived cardiomyocytes on the NEA: (i) Scale bar 200 µm; (ii) Scale bar 50 µm. (c) Quantified viability comparing unmodified and modified NEAs. (n = 3 devices per condition). Error bar represents standard deviation. No significant difference was observed (one-way ANOVA); Schematic representation of the physiological mechanisms underlying observed potential changes: (d) Caffeine-induced calcium release pathway; g) Ouabain-induced Na+/K+-ATPase inhibition pathway; Local K flux recordings showing pharmacological modulation: (e) Baseline, control and Response to caffeine administration; (h) Response to ouabain administration; Quantified drug-evoked potential shifts six recording trials per condition; comparison between control and drug conditions showed statistically significant differences at the 0.05 level (one-way ANOVA, p < 0.05) for caffeine (f) and ouabain (i).

To test KINESIS in a pharmacologically responsive model, we first applied caffeine (10 mM), which induces calcium release and secondary K⁺ efflux via SK channels. Baseline recordings were obtained on day 4. Media-only controls showed minor fluctuations, while caffeine triggered a +11.93 mV shift over 10–20 minutes **(Figure 4d–f)**. This increase reflects localized K⁺ accumulation due to activated efflux, consistent with expected RyR2–SK signaling^33–35^. The shift was significant across replicates **(Figure 4f)**, confirming KINESIS detection of dynamic chemical activity.

To test suppression of K⁺ uptake, we next applied ouabain (0.2 µM), an Na⁺/K⁺-ATPase inhibitor. As expected, ouabain produced a –14.25 mV drop **(Figure 4g–i)**, likely reflecting reduced active K⁺ reuptake and localized depletion at the membrane interface^36,37^. Control DI water produced negligible change. Though bulk K⁺ levels may rise, KINESIS resolves cell-proximal gradients not captured by field-potential approaches.

Both responses stabilized within 5–10 minutes, a timescale consistent with membrane diffusion kinetics and evolving local gradients. Together, these results demonstrate that KINESIS enables label-free, chemically selective, and spatially localized recording of drug-induced potassium flux, establishing its utility for mechanistic screening and cardiotoxicity studies.

## CONCLUSION

KINESIS represents, to date, the first nanoscale ion-selective nanoelectrode array that enables label-free, noninvasive, single-cell measurement of potassium (K⁺) dynamics in live human cardiomyocytes. High-aspect-ratio platinum nanopillars coated with a valinomycin-based membrane create a stable, membrane-proximal interface for localized potentiometric readout.

The platform exhibits near-Nernstian sensitivity and pronounced selectivity versus Na⁺ (KKNa ≪1), delivering stable and reproducible signals suitable for continuous measurements. In pharmacological tests, caffeine elicited a positive potential shift consistent with K⁺ efflux, whereas ouabain produced a negative shift consistent with suppressed K⁺ uptake, confirming functional specificity and physiological relevance. By coupling subcellular access with chemical selectivity, KINESIS addresses a key gap left by conventional NEAs, ISEs, and optical probes, and supports applications in cardiotoxicity screening, ion-channel mechanism studies, and patient-specific disease modeling. The current temporal resolution is limited by diffusion within the ion-selective membrane, which may be improved through thickness optimization and the incorporation of more lipophilic borate additives such as potassium (or sodium) tetrakis[3,5-bis(trifluoromethyl)-phenyl] borate to lower membrane resistance, enhance response speed and sensitivity. Future work will focus on accelerating response kinetics, multiplexing KINESIS for multi-ion detection, and extending the platform to 3D tissue models for greater translational relevance.

## MATERIALS AND METHODS

### Fabrication of NEA

**Step 1: Substrate Preparation and Initial Patterning:** Fused silica wafers were cleaned, primed with adhesion promoter, and coated with AZ1512 positive photoresist. Circular openings were patterned using a maskless aligner (Heidelberg MLA150) and developed. A 240 nm Cr hard mask was deposited by e-beam evaporation and lifted off to define circular disks. **Step 2: Nanopillar Formation and Electrode Lead Patterning:** Quartz nanopillars (∼7 µm height) were etched using the Cr mask in an ICP-RIE system in two steps with intermediate chamber cleaning. Pillar taper was adjusted by wet etching in 20:1 buffered oxide etch (BOE). The Cr mask was removed, and Ti/Pt electrode leads (10 nm/40 nm) were defined by photolithography, sputter deposition, and lift-off. **Step 3: Electrode Metallization and Passivation**: Following lead formation, a passivation stack of SiO₂/SiNₓ was deposited by PECVD at 350 °C. **Step 4: Contact Area Opening and Device Assembly**: Contact areas over nanopillars were opened by photolithography and BOE etching. Wafers were diced, and PDMS wells were attached by oxygen plasma bonding.

### Impedance measurements

Impedance was measured with the MEA-IT system (Multi Channel Systems MCS GmbH) using its default 1 kHz, 100 mV rms sinusoidal excitation. This single-frequency magnitude is the manufacturer’s standard diagnostic output and a field-standard metric for assessing electrode contact quality within the electrophysiological frequency band^24^. Values were averaged across electrodes to monitor relative changes in coupling during culture; no equivalent-circuit analysis or modeling was applied.

### Electrodeposition of Conductive Intermediates

To establish a robust protocol for NEA functionalization, optimization of the intermediate conductive layer and potassium ion-selective membrane (KISM) was first performed on bulk Pt wires. The KSM was prepared by combining valinomycin, potassium tetrakis(p-chlorophenyl)borate (KTpClPB), and dioctyl sebacate (DOS) in tetrahydrofuran (THF) and stirred to ensure complete dissolution. Pt wires were coated with conductive polymers to improve membrane adhesion and charge-transfer kinetics, with PANI:DBSA selected as the primary formulation. PANI:DBSA was electrodeposited, followed by repeated dip-coating cycles in the KISM solution to achieve uniform coverage. Coated electrodes were dried and conditioned in **10⁻⁶ M** KCl overnight (∼12 h) to hydrate the membrane and activate ion exchange sites before calibration. Membrane performance was assessed using a two-electrode potentiometric setup (Figure S3b) to compare two membrane formulations (KSM-1, KSM-2) and evaluate conductive coatings including PPy, PEDOT, unmodified PANI, and PANI composites with graphene oxide or modified dopant content. Results from this screening informed the final membrane–intermediate layer combination used for NEA devices.

Two membrane formulations, denoted KSM-1 and KSM-2, were compared to evaluate the effect of plasticizer-to-ionophore ratios on membrane selectivity and stability as shown in figure S3c. Furthermore, various electrode coatings were systematically tested, including polypyrrole (PPy), poly(3,4-ethylenedioxythiophene) (PEDOT), unmodified PANI, and several PANI: DBSA composites. The PANI composites were further functionalized with graphene oxide (GO) at concentrations of 0.1 and 0.3 mg/mL. In addition, the impact of increased dopant concentration (PANI:3.0DBSA) and PtB deposition were assessed. This comparative analysis provided a framework for identifying material combinations with optimal compatibility, mechanical stability, and electrochemical performance in the context of valinomycin-based potassium sensing. The findings from this phase of optimization directly informed the subsequent integration into the NEA-based sensing platform.

### Ion-Selective Membrane (KSM) Preparation

Two potassium ion-selective membrane (K-ISM) formulations, KSM-1 and KSM-2, were prepared to compare sensing performance and optimize composition. Both used polyvinyl chloride (PVC) as the polymer matrix and dioctyl sebacate (DOS) as a plasticizer in a 1:2 PVC:DOS mass ratio, with valinomycin as the potassium-selective ionophore and potassium tetrakis(4-chlorophenyl)borate (KTClPB) as a lipophilic ionic additive. All components were dissolved in 1 ml tetrahydrofuran (THF) to form homogeneous casting solutions. The specific compositions were as follows: KSM-1: PVC 33.00 mg mL⁻¹, DOS 65.00 mg mL⁻¹, KTClPB 0.53 mg mL⁻¹, and valinomycin 2.05 mg mL⁻¹; KSM-2: PVC 33.00 mg mL⁻¹, DOS 66.70 mg mL⁻¹, KTClPB 1.00 mg mL⁻¹, and valinomycin 1.00 mg mL⁻¹. KSM-1 contained a lower proportion of ionophore and additive relative to plasticizer, while KSM-2 used a higher ionophore and additive content, maintaining the same PVC concentration.

### KSM Functionalization on Bulk Pt Wires

Platinum wires (0.25 mm diameter) were initially cleaned and coated with a conductive polyaniline layer doped with 1.5 dodecylbenzenesulfonic acid (PANI:1.5DBSA) using electrodeposition in order to enhance adhesion of the membrane and facilitate ion-to-electron transduction. The pre-coated wires were then functionalized with the KSM by dipping into the membrane cocktail for 15 seconds, followed by air drying for one minute. This cycle was repeated three times to ensure uniform membrane coverage. After coating, the membranes were conditioned in 10⁻⁶ M KCl solution overnight to equilibrate ion-exchange sites and activate the valinomycin ionophores. This low-activity preconditioning step facilitates equilibration of ionophore–ligand complexes and stabilizes the membrane response, minimizing ion overloading and flux effects that can lead to super-Nernstian behavior and increased detection limits^38^.

### KSM Functionalization of NEAs in KINESIS Platform

To form the intermediate layer, Pt black and PANI:1.5DBSA were coated onto the nanopillars through electrodeposition as shown in Figure S6. A 2 µL of optimized KSM1 was dispensed onto the centre of the device, and immediately (∼1 s) re-aspirated, leaving a thin residual film over the Pt-black-coated nanopillars. The film was cured for 24 h under ambient conditions (20–25 °C, 30–60 % RH). The resulting membrane showed an average thickness of 1.62 ± 0.17 µm (DektakXT Stylus Profiler, Bruker), confirming uniform coverage. Then we conditioned the devices in 10^-6^M KCl. Now the KINESIS are ready for testing. Potentiometric measurements were carried out using a PalmSens4 benchtop potentiostat (PalmSens BV, Houten, NL) operated with PSTrace software (v 5.8). The device was set to open circuit potentiometry mode with a 1 s sampling interval and a total acquisition time of 8000 s per calibration run. Prior to cell experiments, fully functionalized KINESIS devices (PtB + KSM 1) were calibrated potentiometrically in KCl solutions ranging from 1 × 10⁻⁶ to 1 × 10⁻¹ M at 25 °C. Open circuit potential (OCP) was recorded for ≥ 120 s at each concentration.

### Membrane thickness characterization

The final thickness of the potassium-selective membrane was determined using a DektakXT Stylus Profiler (Bruker). Profilometric scans performed across multiple regions of the coated nanopillar array showed a consistent average thickness of 1.62 ± 0.17 µm, confirming the formation of a uniform thin film.

### KINESIS Device Sterilization and Preparation for Cell Culturing

KINESIS devices were sterilized and prepared for cell culture beginning with ultraviolet–ozone (UVO) treatment for 10 min (UVO-CLEANER, Model 42, Jelight Company, Inc.). NEAs were then rinsed three times with phosphate-buffered saline (PBS) followed by two washes with 70% ethanol inside a biosafety cabinet (BSC). Devices were air-dried for 30–60 min and sterilized under UV light in the BSC for 1 h. Following sterilization, NEAs were coated with fibronectin (Millipore Sigma) at 50 µg/ml. An 8 µL droplet of the diluted fibronectin was applied to the center of the NEA, covering the electrode area, and incubated at 37 °C for at least 1 h.

### Cell Seeding and Maintenance

Human iPSC-derived ventricular cardiomyocytes (Celo.Cardiomyocytes, Celogics) were thawed and prepared according to the manufacturer’s instructions. These cells, differentiated from a proprietary human iPSC line (fibroblast origin, Caucasian male donor), exhibit spontaneous beating within 2 days post-thaw and stabilize at 45–60 bpm by day 7. Cultures form synchronous monolayers expressing ventricular markers such as connexin 43, indicative of high interconnectivity. Cells were seeded onto fibronectin-coated NEAs at 50,000 cells in 8 µL and incubated at 37 °C for 60–90 min before the gradual addition of 400 µL plating medium (Celogics) with the NEAs positioned at a 30° angle. NEAs were then returned to a flat position and incubated for 24 h before replacing the plating medium with advanced medium (Celogics). Media changes (50%) were performed every 48 h. By day 5, cultures were used for electrophysiology experiments. Following experiments, NEAs were cleaned for reuse by incubation in enzymatic cleaner (Boston protein remover) for 12–14 h, followed by immersion in a 1% Terg-A-Zyme solution (Sigma) in distilled water overnight at room temperature with gentle shaking. NEAs were rinsed thoroughly with distilled water before reuse.

### Pharmacological Experiments

Pharmacological experiments were carried out using ouabain and caffeine. For the ouabain experiments, deionized (DI) water was used as the control, and 2 µL was added. Ouabain was dissolved in DI water to prepare a 1 M stock solution. For the caffeine experiments, cell culture media was used as the control. Caffeine was dissolved in media to prepare a 77 mM stock solution.

### Cell Viability Assessment on Nanoelectrode Arrays

Cell viability on unmodified and surface modified nanoelectrode arrays was measured using a standard fluorescence based LIVE DEAD assay following the manufacturer’s protocol (Thermo Fisher Scientific). Cardiomyocytes cultured on the devices were gently rinsed with warm RSPBS, incubated in the staining solution for approximately 30 min, washed, and imaged immediately at 10x magnification. Multiple non-overlapping fields were collected per device under identical acquisition settings. Viability was quantified using image-based analysis in ImageJ, and the percent viable cells was averaged across replicate devices for each condition.

## SUPPORTING INFORMATION

The following files are available free of charge.

- Detailed fabrication workflow and optimization steps for potassium-selective nanoelectrode arrays (NEAs), including nanopillar array preparation, electrodeposition of intermediate conductive layers, and ion-selective membrane functionalization. Additional data include Nernstian calibration plots, K⁺/Na⁺ selectivity measurements, cyclic voltammetry and impedance characterization, SEM imaging of electrode structures, and real-time potentiometric recordings from single human iPSC-derived cardiomyocytes under pharmacological modulation (PDF).
- Illustration of monolayer confluency and spatial distribution of hiPSC-CMs across the platform (MP4).

## AUTHOR INFORMATION

### Corresponding Author

Zeinab Jahed, PhD – Aiiso Yufeng Li Family Department of Chemical and Nano Engineering, University of California San Diego, La Jolla, CA 92093, USA; Shu Chien-Gene Lay Department of Bioengineering, University of California San Diego, La Jolla, CA 92093, USA; Joseph Wang – Aiiso Yufeng Li Family Department of Chemical and Nano Engineering, University of California San Diego, La Jolla, CA 92093, USA;

### Notes

The authors declare no competing financial interest.

## ACKNOWLEDGMENTS

This work was in part supported by the Air Force Office of Scientific Research YIP award (AFOSR FA9550-23-1-0090) and UC San Diego Materials Research Science and Engineering Center (UC San Diego MRSEC), supported by the National Science Foundation (Grant DMR-2011924) to Z.J.

## ABBREVIATIONS

ISE: ion-selective electrode
ISM: ion-selective membrane
KINESIS: K⁺-Ion Nano-Electrode Selective Interface System
KSM: potassium ion-selective membrane
KTpClPB: potassium tetrakis(4-chlorophenyl)borate
MEA: microelectrode array
NEA: nanoelectrode array
OCP: open-circuit potential
PANI:1.5DBSA: polyaniline doped with 1.5 equiv dodecylbenzenesulfonic acid
RyR2: ryanodine receptor 2
THF: tetrahydrofuran
hiPSC-CM: human induced pluripotent stem cell–derived cardiomyocyte

