## Supplementary info for "Potassium-Selective Nanoelectrode Arrays for Single-Cell Profiling of human iPSC-Derived Cardiomyocytes"

### Supplementary Information

*Dhivya Pushpa Meganathan*[1], *Romeo Banzon*[1], *Ana Casanova*[1], *Einollah Sarikhani*[1],*Kuldeep Mahato*[1] *Hillary Vu*[1], *Sarah Reade*[1], *Iswerya Ambika Devarajan*[1], *Anum Tahir*[1,3], *Lekshmi Sasi*[1], *Leah Sadr*[1], *Joseph Wang*[1], *Zeinab Jahed*[1,2]

[1]Aiiso Yufeng Li Family Department of Chemical and Nano Engineering, University of California San Diego, La Jolla, San Diego, CA 92093, USA

[2]Shu Chien-Gene Lay Department of Bioengineering, University of California San Diego La Jolla, San Diego, CA 92093, USA

[3] Department of Chemistry and Biochemistry, Brigham Young University, Provo, UT, 84602, USA

Corresponding Authors:

Joseph Wang, Zeinab Jahed, PhD

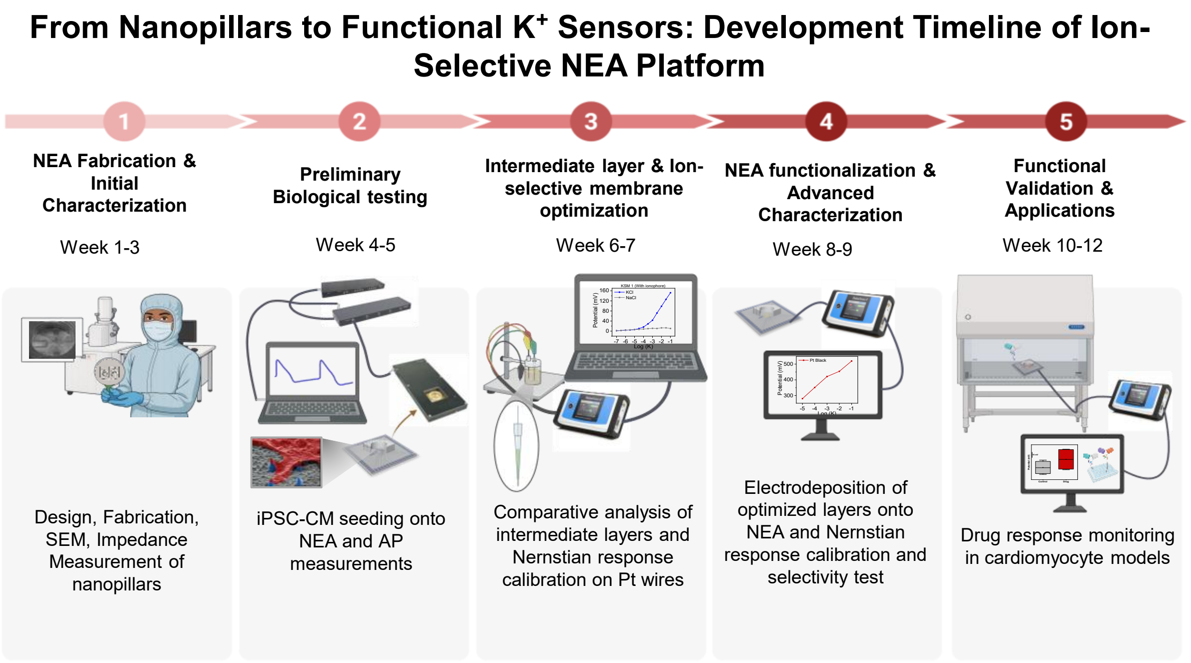


This figure illustrates the systematic development pathway of our ion-selective nanoelectrode array (NEA) platform optimized for potassium (K+) detection. The timeline spans 12 weeks divided into five sequential phases, each representing critical developmental milestones:

**Phase 1 (Week 1-3):** NEA Fabrication and Initial Characterization The foundation of the platform begins with nanopillar design, fabrication using advanced nanofabrication techniques, and initial structural characterization via scanning electron microscopy (SEM). Baseline electrical properties were established through impedance measurements of the bare nanopillars.

**Phase 2 (Week 4-5):** Preliminary Biological Testing Following fabrication, induced pluripotent stem cell-derived cardiomyocytes (iPSC-CMs) were seeded onto the NEA platform to evaluate biocompatibility and establish baseline action potential (AP) measurements. This phase validated the platform's capability for cellular interfacing without significant cytotoxicity.

**Phase 3 (Week 6-7):** Intermediate Layer and Ion-Selective Membrane Optimization This critical phase focused on comparative analysis of various intermediate layer compositions and their integration with the nanoelectrodes. Nernstian response calibration was performed using platinum wire references to optimize selectivity coefficients for potassium ions against potential interferents.

**Phase 4 (Week 8-9):** NEA Functionalization and Advanced Characterization Optimized ion-selective layers were electrodeposited onto the NEA surface, followed by comprehensive characterization of membrane integrity. Nernstian response calibration was conducted and selectivity tests were performed against common interfering ions (Na+) to validate K+ specificity.

**Phase 5 (Week 10-12):** Functional Validation and Applications The fully developed K+-selective NEA platform underwent rigorous testing in cardiomyocyte models to monitor drug-induced changes in potassium dynamics. Real-time monitoring capabilities were demonstrated under various pharmacological interventions that modulate potassium channel activity.

This developmental pipeline represents a significant advancement in nanoscale biosensing technology, enabling selective monitoring of K+ flux

**Figure S1:** Development Timeline of Ion-Selective NEA Platform

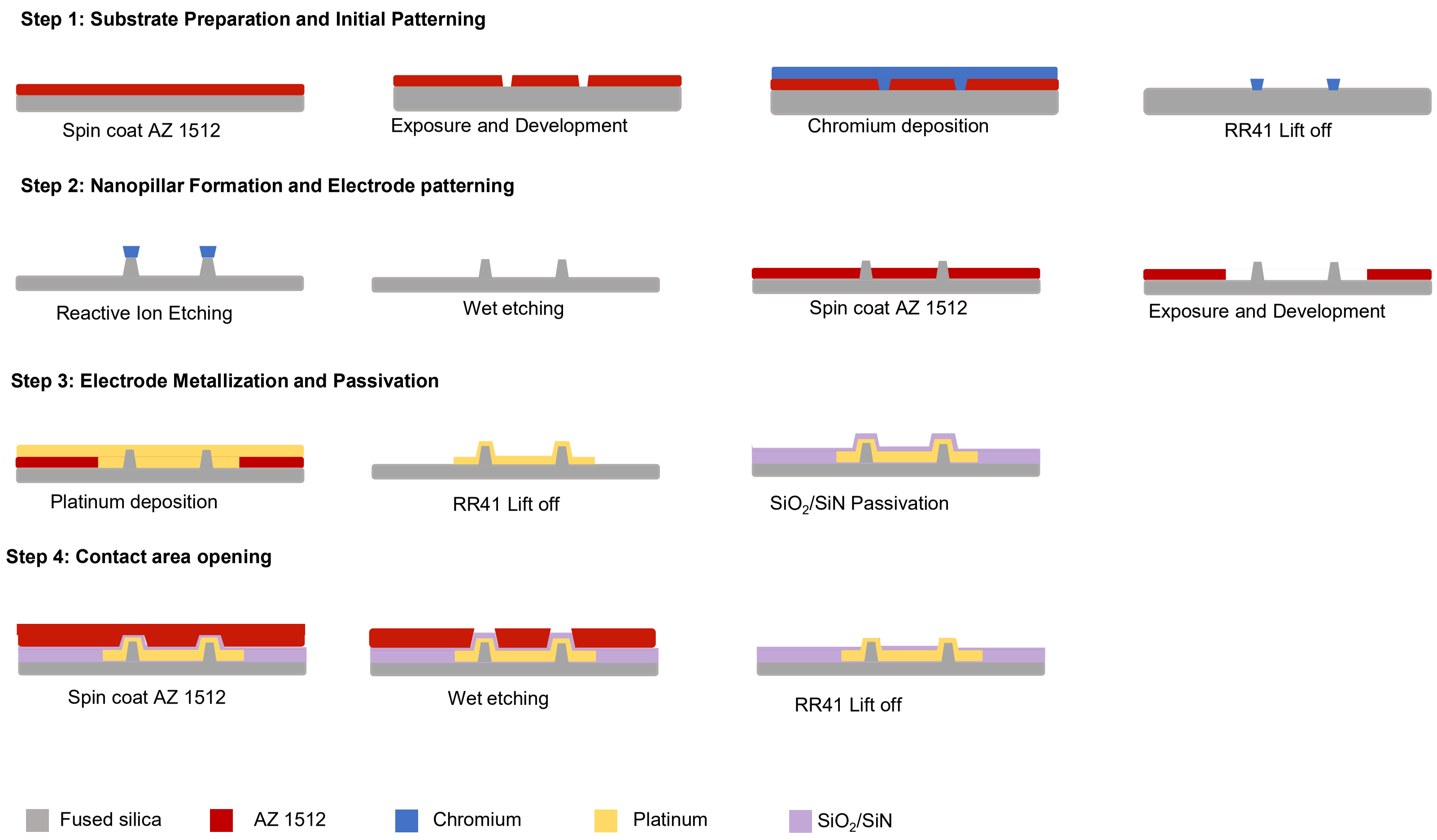


**Figure S2.** Schematic illustration of the nanopillar electrode array fabrication process.

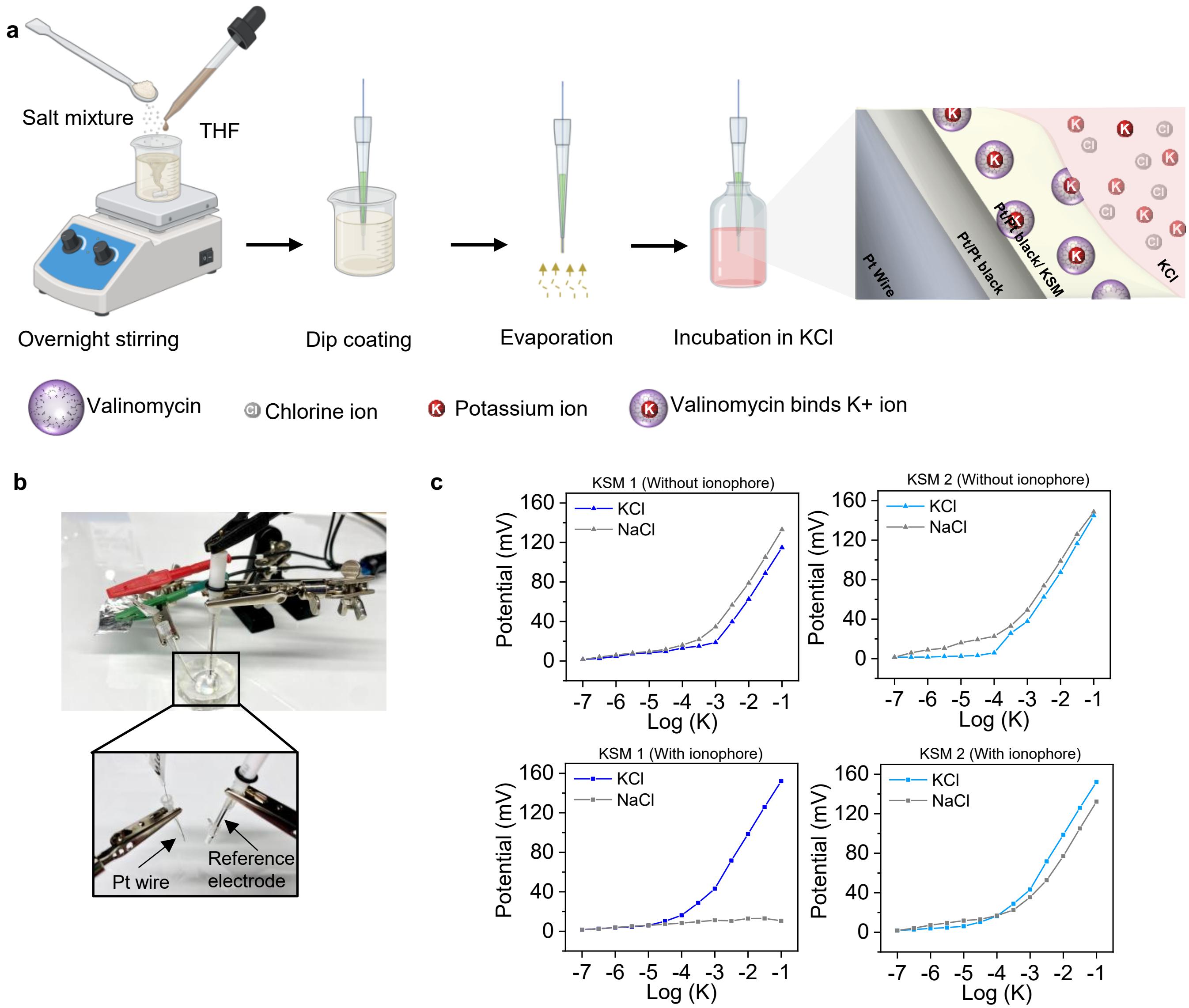


**Figure S3.** Optimization of Potassium Ion-Selective Membranes. (a) Schematic of the Pt wire functionalization process. (b) Photograph of the two-electrode potentiometry setup. (c) Optimization of potassium ion-selective membranes (KSM1 and KSM2) with ionophore and without ionophore, showing sensitivity and selectivity against NaCl.

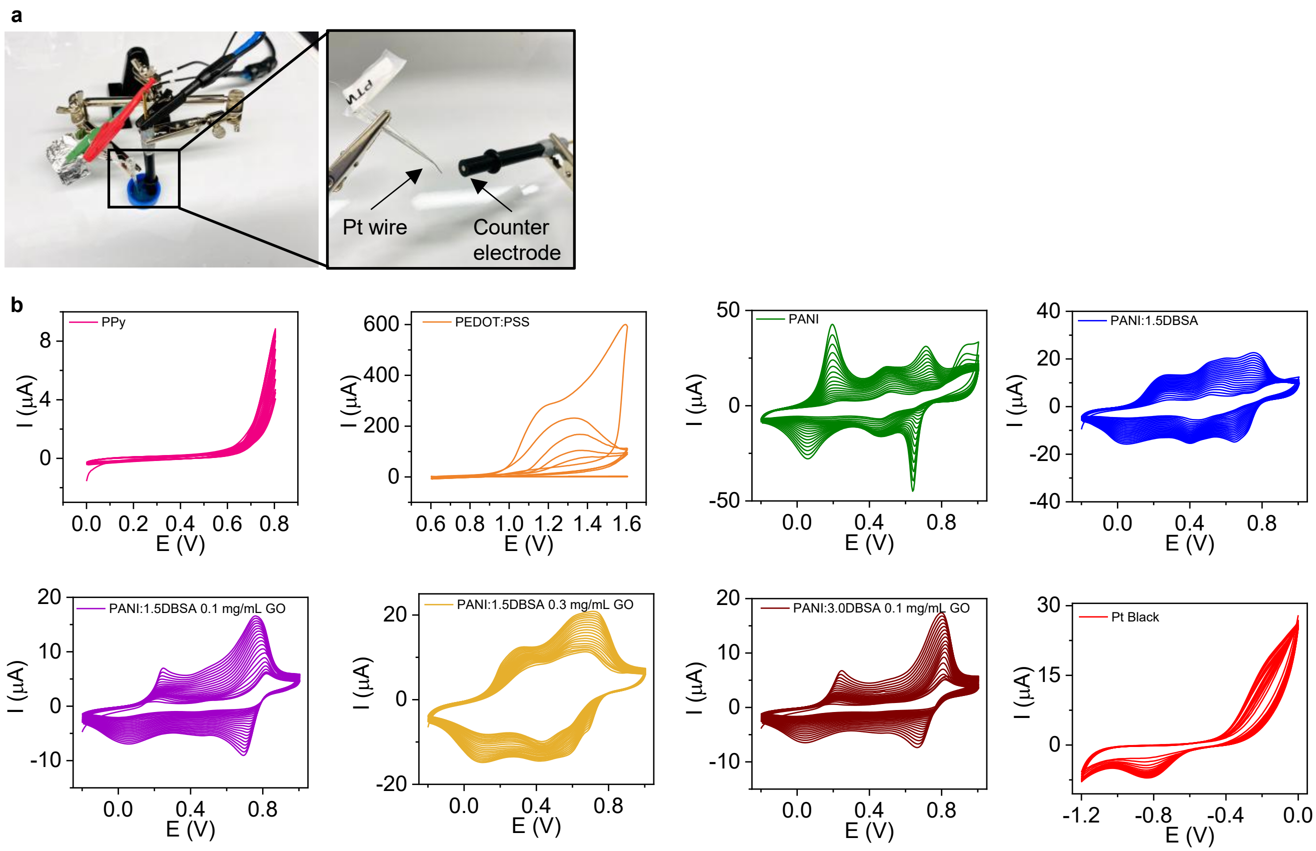


**Figure S4.** Electrodeposition of Conductive Materials on Platinum Wire and Cyclic Voltammetry Characterization. (a) Photograph of the two-electrode electrodeposition setup for coating platinum wire with conductive materials. Inset shows magnified view of the counter electrode and Pt wire configuration during the electrodeposition process. (b) Cyclic voltammograms of various conductive materials electrodeposited on Pt wire: (i) PPy; (ii) PEDOT; (iii) PANI; (iv) PANI:1.5DBSA; (v) PANI:1.5DBSA with 0.1 mg/mL GO; (vi) PANI:1.5DBSA with 0.3 mg/mL GO; (vii) PANI:3.0DBSA with 0.1 mg/mL GO; (viii) Pt Black.

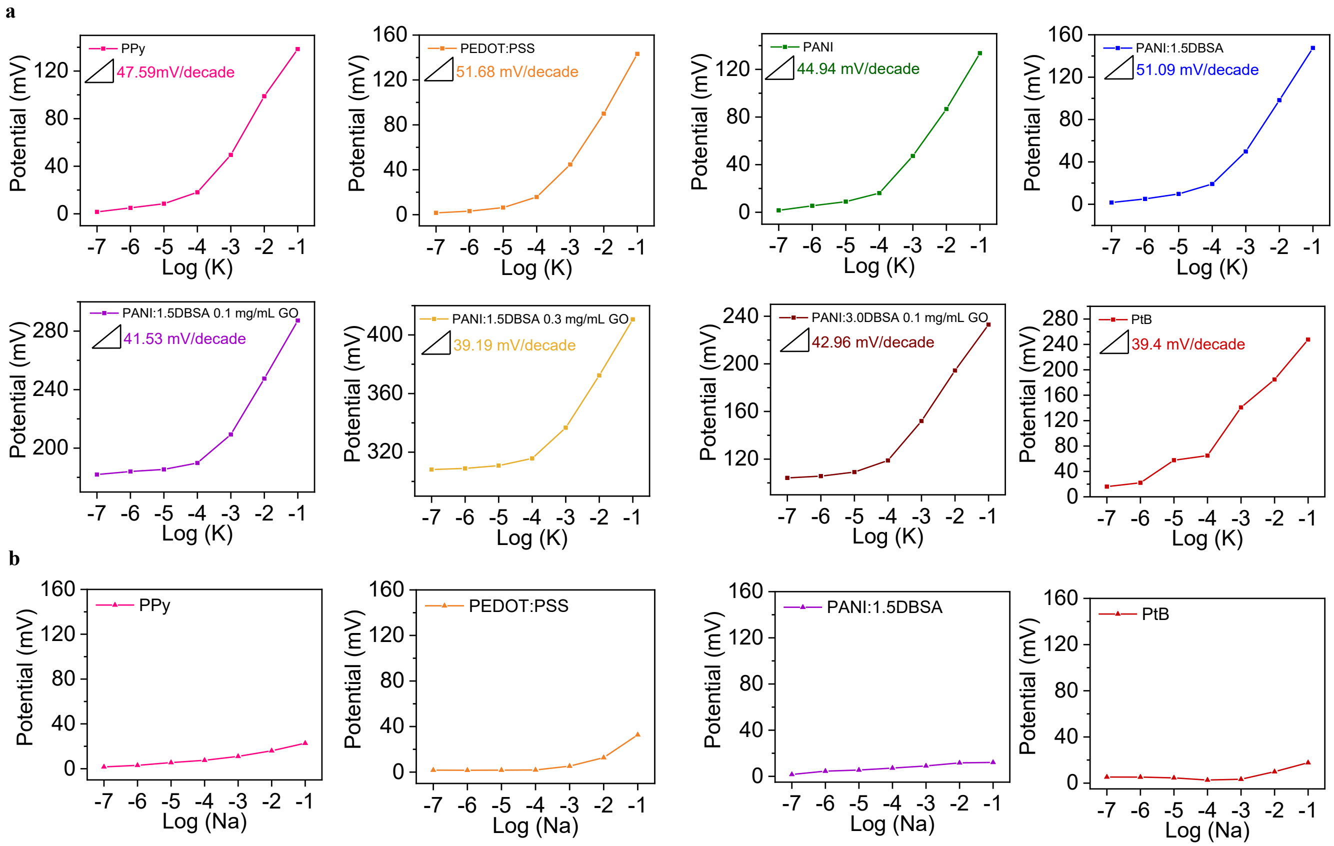


**Figure S5:** Comparative Analysis of Intermediate Conductive Layers for K+ Detection. (a) Sensitivity comparison of various intermediate layers on Pt wires for K+ detection: (i) Polypyrole; (ii) PEDOT; (iii) PANI; (iv) PANI:1.5DBSA; (v) PANI:1.5DBSA with 0.1 mg/mL GO; (vi) PANI:1.5DBSA with 0.3 mg/mL GO; (vii) PANI:3.0DBSA with 0.1 mg/mL GO; (viii) Pt Black. (b) Selectivity comparison of intermediate layers on Pt wires against NaCl interference: (i) Polypyrole; (ii) PEDOT; (iii) PANI:1.5DBSA; (iv) Pt Black.

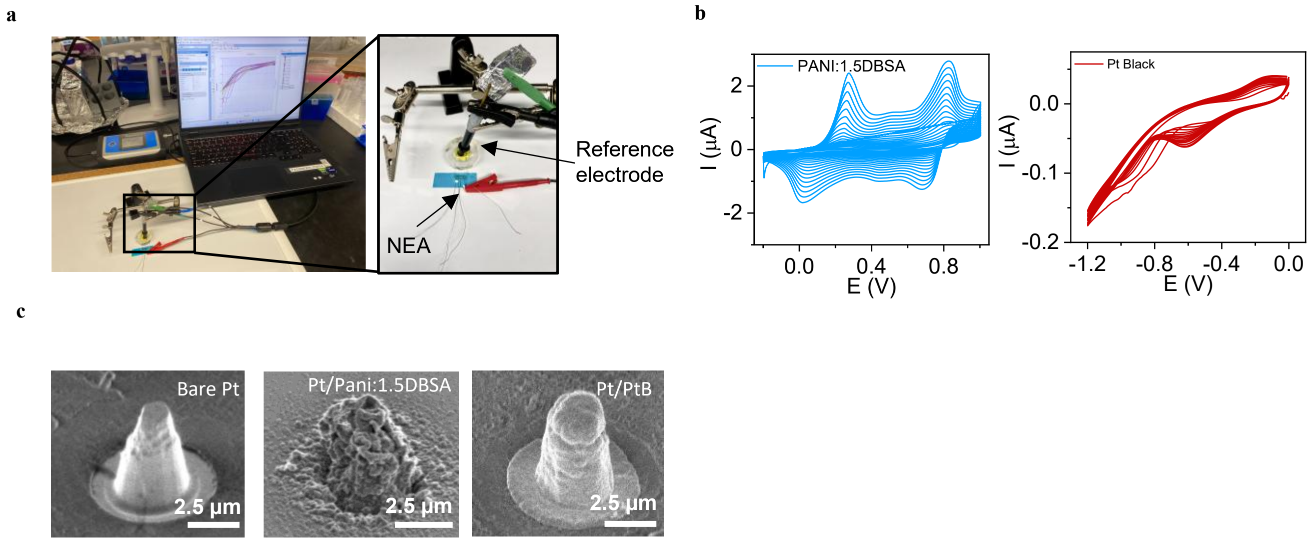


**Figure S6.** Electrodeposition of Conductive Materials on Nanoelectrode Array (NEA) and Cyclic Voltammetry Characterization. (a) Photograph of 8-channel nanoelectrode array (NEA). (b) Photograph of the two-electrode electrodeposition setup for coating platinum nanopillar with conductive materials, showing the computer interface, potentiostat, and electrode arrangement. Inset shows magnified view of the electrochemical cell configuration, identifying the reference electrode and NEA positions. (c) Cyclic voltammograms of various conductive materials electrodeposited on nanopillars: (i) PANI:1.5DBSA; (ii) Pt Black. (d) SEM images of Pt nanopillars: (i) Before electrodeposition; (ii) After electrodeposition of PANI:1.5DBSA; (iii) After electrodeposition of Pt Black. (Scale bar: 2.5 µm)

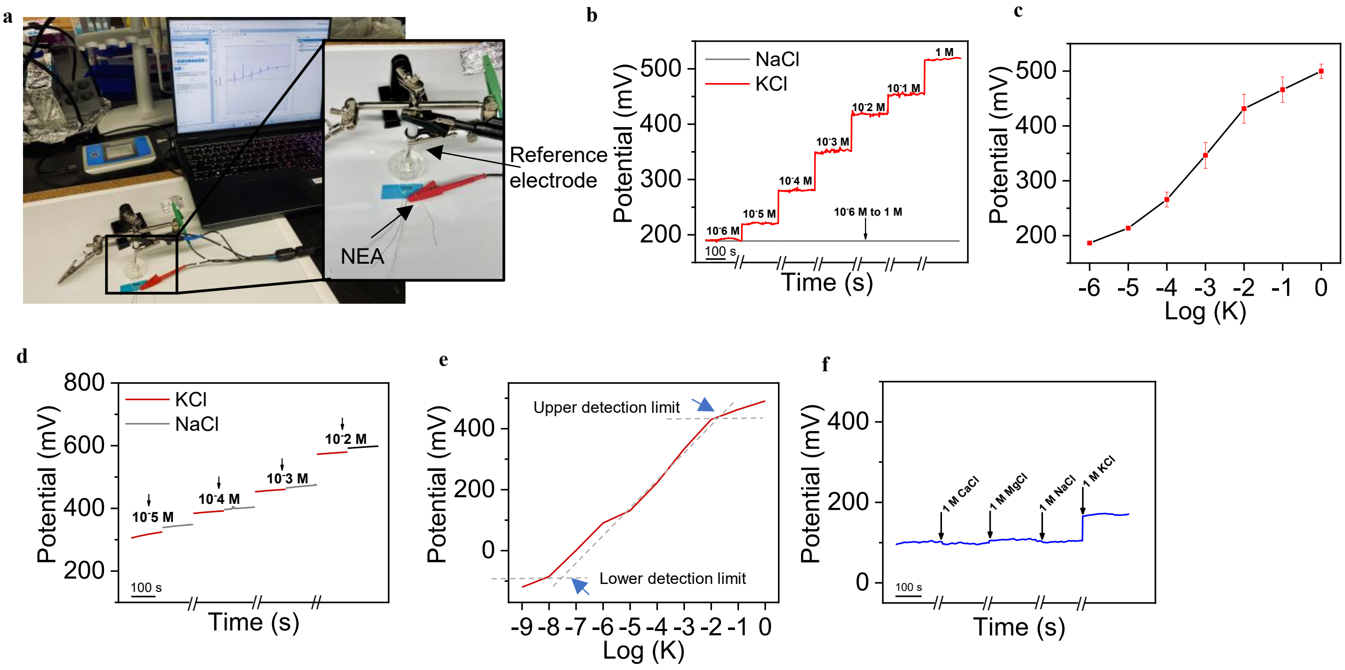


**Figure S7.** Electrochemical Characterization of fully functionalized Nanoelectrode Array (NEA) (a) Photographic image of the potentiometric measurement setup showing the NEA and reference electrode (b) Time dependent response of potential vs. time showing K+ sensitivity and Selectivity evaluation against NaCl interference (c) Potassium calibration curves (potential vs. log[K+] concentration): Pt Black, showing K+ sensitivity and averaged across 8 measurements across various electrodes (d) Time-dependent potential responses upon sequential addition of KCl (red line) and NaCl (gray line) solutions at increasing concentrations ($10^{-5}$ M to $10^{-2}$ M) (e) Potential response of $K^+$-versus Log[K], indicating upper and lower detection limits. (f) Representative potentiometric trace showing selective response of the $K^+$-selective nanopillar electrode to common physiological ions ($Ca^{2+}$, $Mg^{2+}$, $Na^+$ and $K^+$).

a

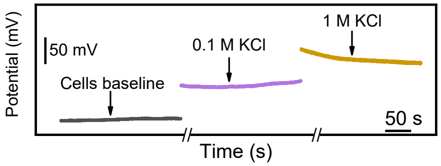


**Figure S8:** (a) Local K+ flux potential recordings showing baseline iPSC-CM activity and response to potassium concentration changes: (i) Addition of 0.1 M KCl; (ii) Addition of 1 M KCl.

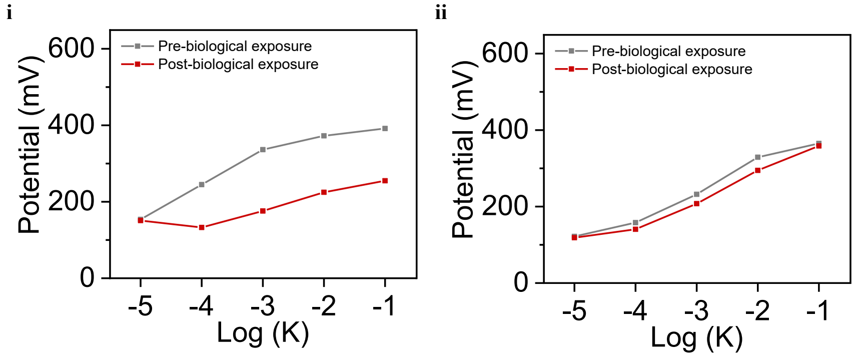


**Figure S9:** (a) Effect of biological exposure on $K^+$ calibration. Potentiometric calibration curves (potential vs log[$K^+$]) for three representative electrodes (i) and (ii), measured before biological exposure (gray) and after two 5-day cell-culture exposures on fibronectin-coated devices (red)

To evaluate robustness under biologically relevant conditions, we compared $K^+$ calibration curves recorded before and after repeated exposure to cell-culture environments. Each device was first calibrated in KCl solutions ($10^{-6}$ M) to establish the pre-biological-exposure response. The same electrodes were then coated with fibronectin and seeded with human iPSC-derived ventricular cardiomyocytes (Celogics) using the manufacturer-supplied plating medium and supplements. Cultures were maintained for five days under standard incubator conditions, with partial medium changes every 48 h.

After each culture period, cells were detached with TrypLE, and the electrodes were cleaned sequentially with enzymatic cleaner (Boston Protein Remover) and detergent (Tergazyme), thoroughly rinsed with DI water, re-conditioned in $10^{-6}$ M KCl for atleast 24 h, and then reused for the next culture. Following the second culture cycle, the same devices were re-calibrated under identical conditions to obtain the post-biological-exposure curves.

In the plot, the pre-biological-exposure data represent the initial calibration of each electrode prior to any cell-culture contact, whereas the post-biological-exposure data correspond to the same electrodes after two full cardiomyocyte culture cycles and prolonged exposure to fibronectin coating, cell attachment, and protein-containing media. Across three representative electrodes, the calibration slopes changed as follows: Figure 3 (f) Electrode 1: 66.5 → 59.3 mV $dec^{-1}$ (–10.9%), Figure S9 (i) Electrode 2: 76.2 → 60.3 mV $dec^{-1}$ (–20.8%), Figure S9 (ii)Electrode 3: 65.7 → 63.4 mV $dec^{-1}$ (–3.5%)
The average slope change was –11.7%, reflecting only modest sensitivity loss consistent with minor fouling or membrane aging, and no catastrophic degradation of ion-to-electron transduction. These results support robust operation of the sensors under the intended in-vitro use conditions.
